# Hepatic JNK-mediated bile acid homeostasis regulates liver cancer through PPARα

**DOI:** 10.1101/783761

**Authors:** Elisa Manieri, Laura Esteban-Lafuente, María Elena Rodríguez, Luis Leiva-Vega, Chaobo Chen, Francisco Javier Cubero, Tamera Barrett, Julie Cavanagh-Kyros, Davide Seruggia, Maria J. Monte, Jose J.G. Marin, Roger J. Davis, Alfonso Mora, Guadalupe Sabio

## Abstract

cJun NH_2_-terminal kinase (JNK) inhibition has been suggested as a potential treatment for insulin resistance and steatosis through activation of the transcription factor PPARα. However, the long-term consequences have not been evaluated. We found that hepatic JNK deficiency alters bile acid and cholesterol metabolism, resulting in hepatic expression of FGF15 and activation of ERK in cholangiocytes, which ultimately promotes their proliferation. Genetic inactivation of PPARα identifies PPARα hyperactivation as the molecular mechanism for these deleterious effects. Our analysis indicates that hepatic PPARα activation is oncogenic: PPARα deficiency protects mice against carcinogen-induced hepatocellular carcinoma under high fat diet (HFD) condition. These surprising results urge the re-consideration of using JNK inhibitors or PPAR agonists for the treatment of metabolic syndrome.

## INTRODUCTION

Liver cancer is the fifth most common cancer and the second leading cause of cancer deaths worldwide (El-Serag, Davila et al., 2003, Parkin, Bray et al., 2005). Metabolic syndrome is a newly recognized, but important risk factor thought to contribute to the increased incidence of hepatocellular carcinoma (HCC) (Klein, Dawson et al., 2014). Steatosis contributes to HCC development due to its association with oxidative stress and inflammation (Smedile & Bugianesi, 2005). Metabolic disorders are common among obese and diabetic patients, and hepatocellular injuries can occur due to increased fat accumulation in the liver. Non-alcoholic fatty liver disease (NAFLD) is extremely frequent in these patients (Caldwell, Crespo et al., 2004), and body weight excess is commonly associated with advanced disease (Neuschwander-Tetri, Brunt et al., 2003). Recent studies have led to the identification of hepatic cJun NH_2_-terminal kinase (JNK) as a signal transduction pathway that is critically required for obesity-induced insulin resistance and steatosis (Manieri & Sabio, 2015). The cJun NH_2_-terminal kinase (JNK) signaling pathway contributes to the development of obesity and insulin resistance (Sabio & Davis, 2010). Indeed, mice deficient for *Jnk* in hepatocytes are resistant to high fat diet (HFD)-induced insulin resistance and steatosis (Vernia, Cavanagh-Kyros et al., 2014); therefore, this signaling pathway represents a potential target for therapeutic intervention. Biochemical studies demonstrate that JNK suppresses PPARα activation in hepatocytes, affecting lipid metabolism and steatosis through the hepatokine FGF21 (encoded by a PPARα target gene) (Vernia, Cavanagh-Kyros et al., 2016, Vernia et al., 2014).

The transcription factor PPARα plays a pivotal role in intracellular free fatty acid (FFA) and triglyceride metabolism by regulating genes involved in fatty acid transport and degradation in mitochondria and peroxisomes (Evans, Barish et al., 2004, Gulick, Cresci et al., 1994, Unger & Zhou, 2001). PPARα is expressed primarily in liver, heart, and muscle and is a major regulator of fatty acid transport, catabolism and energy homeostasis (Memon, Tecott et al., 2000). PPARα activation in the liver is increased in metabolic diseases and obesity (Memon et al., 2000), and PPARα agonists appear to be therapeutically beneficial in diabetes. In fact, PPARα protects against steatosis in the mouse (Ip, Farrell et al., 2003) and suppresses hepatic inflammation (Teoh, Williams et al., 2010). However, PPARα deficiency in mice increases susceptibility to diethylnitrosamine (DEN)-induced HCC (Zhang, Chu et al., 2014), but long-term studies in rodents showed an association of PPARα agonists with hepatic carcinogenesis (Holden & Tugwood, 1999). These findings conflict with the growth inhibitory effects reported for PPARα agonists in cancer cell lines, including HCC cell lines (Maggiora, Oraldi et al., 2010, Panigrahy, Kaipainen et al., 2008, Yamasaki, Kawabe et al., 2011). PPARα may therefore cause context-specific actions on liver cancer development.

The activation of PPARα modifies bile acid (BA) synthesis, conjugation and transport (Li & Chiang, 2009). Altered regulation of BA may protect against steatosis but could increase liver cancer development due to changes in FGF protein levels. Although FGF19 improves the glycemic response and reduces liver steatosis, it also induces liver cancer (Shapiro, Kolodziejczyk et al., 2018). These contrasting potential functions of BAs prompted us to examine whether lack of JNK in hepatocytes, and the resulting hyperactivation of PPARα, could alter BA homeostasis with subsequent deleterious effects. Elucidation of the contribution of JNK and PPARα to liver carcinogenesis may help the development of effective treatments against this malignancy.

## RESULTS

### Hepatic JNK-deficiency alters bile acid homeostasis

We have previously shown that hepatic JNK deficiency results in the activation of the nuclear transcription factor PPARα and protection against diet-induced insulin resistance and steatosis (Vernia et al., 2014). The activation of PPARα caused altered BA metabolism (Li & Chiang, 2009). We therefore examined BA in hepatic JNK deficient mice (L^DKO^) and control mice (L^WT^) at 6 months of age. We found that total BA concentration in the blood of L^DKO^ mice were significantly increased compared with L^WT^ mice (Fig 1A). The increase in circulating BA concentration is consistent with the possibility that L^DKO^ mice may suffer from cholestasis.

**Figure 1.**
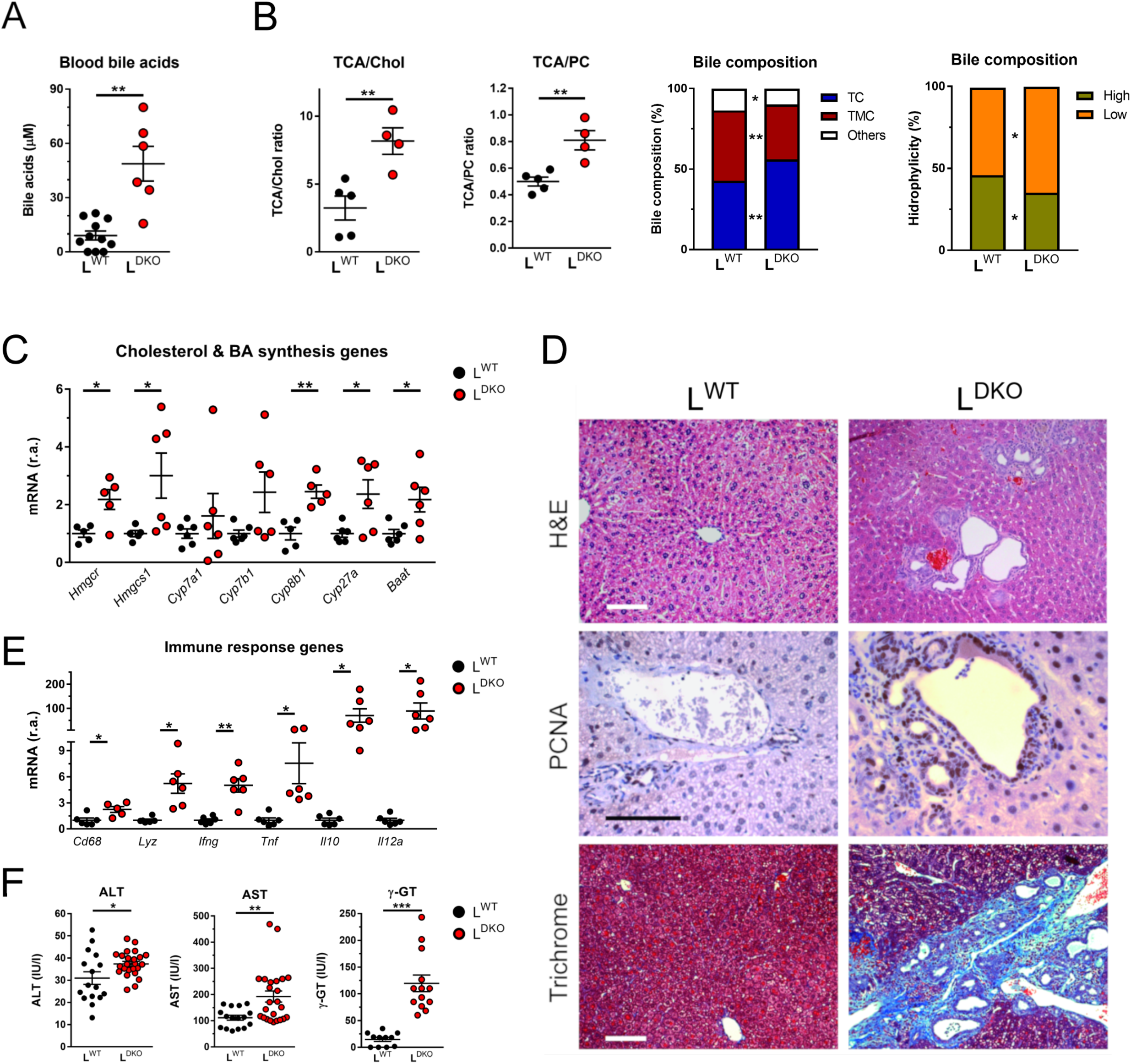
Hepatic JNK-deficiency alters bile acid production and causes the development of cholestasis. A L^WT^ and L^DKO^ mice at 6 months of age were fasted overnight and blood was collected. The amount of bile acids in the blood was measured (mean ± SEM; n = 6-11). Statistically significant differences between L^DKO^ and L^WT^ are indicated (**, *P* < 0.01). B The composition of bile fluid collected from the gall bladder was examined by measurement of the ratio of bile acids (BA) to cholesterol (Chol) or phosphatidylcholine (PC) and the different type of BA. The data presented are the mean ± SEM (n = 4-5). Statistically significant differences between L^DKO^ and L^WT^ are indicated (*, *P* < 0.05; **, *P* < 0.01). C L^KO^ and L^WT^ mice (age 6 months) were fasted overnight prior to removal of the liver. The expression of genes related to cholesterol synthesis (*Hmgcr* and *Hmgcs1*) and bile synthesis (*Cyp7a1*, *Cyp7b1*, *Cyp27a1, Baat*, *Cyp27a*, *Cyp8b1*) was measured by quantitative RT-PCR (mean ± SEM; n = 5-6). The expression was normalized to the amount of *18S* RNA in each sample. Statistically significant differences between LD^KO^ and L^WT^ are indicated (*, P < 0.05; **, P < 0.01). D Representative liver sections stained with hematoxylin and eosin (H&E), an antibody to PCNA, and Masson Trichrome (Trichrome) are presented. Scale bar = 100 µm. E The expression of genes related to inflammation was evaluated by RT-qPCR. (mean ± SEM; n = 5-6). The expression was normalized to the amount of *18S* RNA in each sample. Statistically significant differences between LD^KO^ and L^WT^ are indicated (*, P < 0.05). F Liver damage was assessed from serum measurements of ALT, AST and γ-GT. (mean ± SEM; n = 11-24). Statistically significant differences between LD^KO^ and L^WT^ are indicated (*, P < 0.05; **, P < 0.01, ***, P < 0.001).

The analysis of bile collected from the gallbladder of L^DKO^ and L^WT^ mice revealed significantly increased amounts of BA relative to the amount of cholesterol and phosphatidylcholine (PC) (Fig 1B). Hepatic expression of genes related to hepatic PC synthesis (*Scd2*, *Chpt1*, and *Chkb*) or hepatocyte-mediated transport of PC (*Abcb4* and *Atp8b1*) and BA (*Abc11* and *Slc10a1*) was markedly increased in L^DKO^ mice (Supplementary Fig EV1A, B). Similarly, increased expression of genes related to cholesterol synthesis (*Hmgcs1, Hmgcr*) and BA synthesis (*Baat*, *Cyp8b1 and Cyp27a*) was detected in L^DKO^ mice (Fig 1C). These data are consistent with altered biosynthesis and secretion of both cholesterol and BA through PPARα activation.

### Hepatic JNK-deficiency causes cholestasis and liver damage

It is established that cholangitis is a major risk factor for the development of cholangiocarcinoma (de Groen, Gores et al., 1999). We therefore examined the liver of mature adult L^DKO^ and L^WT^ mice. No evidence of hepatic disease was found in L^WT^ mice. Similarly, analysis of hepatic sections prepared from young adult L^DKO^ mice (age 4 months) did not indicate the presence of liver pathology (Figure EV1C). However, at age 10 months, 82% of L^DKO^ mice displayed multifocal bile duct hyperplasia together with fibrosis and inflammatory cell infiltrates (Fig 1D). Cholangiocytes stained positively with PCNA, a marker for proliferation (Fig 1D). These changes were associated with increased expression of myeloid genes (*Cd68* and *Lyz*) and inflammatory cytokines (*Ifng*, *Tnf*, *Il10*, and *Il12a*) in the liver (Fig 1E), and increased liver damage, as suggested by the high levels of liver enzymes (ALT, AST, and γ-GT) in the blood of L^DKO^ mice (Fig 1F). Theremaining L^DKO^ mice exhibited cholangiocarcinoma (6%) or appeared to be healthy (12%). At age 14 months, 95% of L^DKO^ mice displayed cholangiocarcinoma (Fig 2A) associated with fibrosis (Fig 2B) and a large increase in liver mass together with a significant increase in ALT and AST (Fig 2C). The remaining L^DKO^ mice (6%) exhibited cystic livers with bile duct hyperplasia. Histological analysis indicated increased staining of glutamine synthetase (GS) in liver tumor lesions, together with neoplastic nodules with positive staining of the ductular markers CK19 and Sox9 (Fig 2D). Together, these data confirm that the majority of mature mice with compound deficiency of JNK1 and JNK2 progressively develop cholangiocarcinoma.

**Figure 2.**
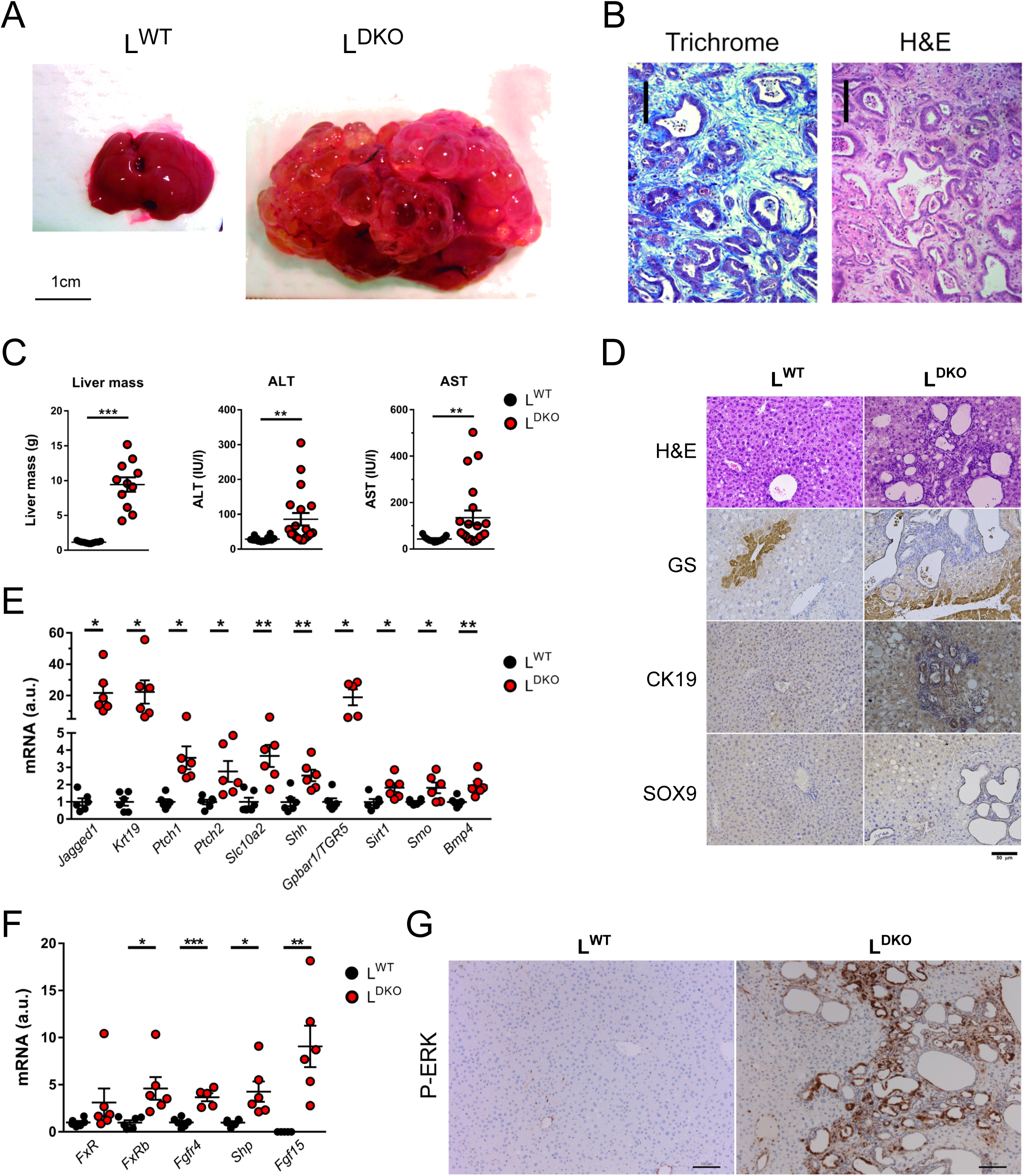
Hepatic JNK-deficiency progress to cholangiocarcinoma through ERK activation. A Representative livers of L^WT^ and L^DKO^ mice at age 14 months are shown. B Representative sections of the liver of 14 month old chow-fed L^DKO^ mice were stained with hematoxylin and eosin (H&E) and Masson Trichrome (Trichrome). Scale bar = 100 µm. C The liver mass and liver damage measured by levels of ALT and AST (mean ± SEM; n = 11-20) is presented. Statistically significant differences between LD^KO^ and L^WT^ are indicated (**, P < 0.01, ***, P < 0.001). D Representative liver sections of 10 months old L^DKO^ and L^WT^ mice stained with glutamine synthetase (GS), Cytokeratin 19 (CK19) and Sox9. Scale bar = 50 µm. E The expression of genes related to cholangiocytes proliferation was evaluated by RT-qPCR in 14-month-old L^WT^ and L^DKO^ mice (mean ± SEM; n = 5-6). Statistically significant differences between LD^KO^ and L^WT^ are indicated (*, P < 0.05; **, P < 0.01). F The expression of genes related to the nuclear factor FXR (*Fxr*, *Fxrb*, *Shp, Fgr4*) and *Fgf15* was measured by quantitative RT-PCR in L^WT^ and L^DKO^ liver from mice at age 6 months (mean ± SEM; n = 5-6). Statistically significant differences between LD^KO^ and L^WT^ are indicated (*, P < 0.05; **, P < 0.01, ***, P < 0.001). G Representative liver sections of 10-month-old L^DKO^ and L^WT^ mice stained with Phospho-ERK. Scale bar = 100 µm.

The development of bile duct hyperplasia and cholangiocarcinoma in L^DKO^ mice was associated with increased hepatic expression of *Cytokeratin 19* (*Krt19*), a cholangiocyte-specific epithelial marker, the G protein-coupled BA receptor 1 (*Gpbar1*) and the apical sodium-dependent BA transporter (Asbt, gene symbol *Slc10a2*) both expressed in cholangiocytes (Keitel, Reinehr et al., 2007) (Dawson, Lan et al., 2009) (Figure 2E). The Notch receptor ligand *Jagged-1* promotes the formation of intrahepatic bile ducts (Piccoli & Spinner, 2001), and was overexpressed in the liver of L^DKO^ mice (Fig 2E). Moreover, Bone morphogenetic protein 4 (Bmp4) mediates cholestasis-induced fibrosis (Fan, Shen et al., 2006) and cooperates with FGF to promote the development of cholangiocytes from hepatoblasts (Yanai, Tatsumi et al., 2008); expression of hepatic *Bmp4* was increased in L^DKO^ mice compared with L^WT^ mice (Fig 2E). Hepatoblasts maturate to cholangiocytes through activation of ERK pathway (Yang, Wang et al., 2017) and BA can increase proliferation by ERK activation through FXR/FGF15/FGR4 pathway (Li & Chiang, 2015). We evaluated this pathway in 6 months-old mice, before cancer has developed. In concordance with elevated BA production, we found high FXR activation as suggested by the high levels of its target genes (*Shp* and *Fgr4*) observed in L^DKO^ mice (Fig 2F). Moreover, while control mice did not expressed *Fgf15*, we could detect *Fgf15* in L^DKO^ livers (Fig 2F). In agreement with these results, histological analysis indicated increased staining of ERK phosphorylation in cholangiocytes from L^DKO^ mice compared with L^WT^ mice (Fig 2G). Together, these changes in FXR/FGF15/FGR4/ERK pathway may contribute to cholangiocyte maturation and proliferation from hepatoblast resulting in bile duct hyperplasia and development of cholangiocarcinoma detected in L^DKO^ mice.

### PPARα deficiency reduces liver cancer in JNK1/2 deficient liver

To confirm that hyperactivation of PPARα is involved in the development of cholangiocarcinoma in L^DKO^ mice, we ablated the *Ppara* gene in L^DKO^ mice. In the resulting JNK1/2 plus PPARα liver-deficient-mice (L^PPARαDKO^), both tumor burden and incidence were clearly reduced (Fig 3A) compared with L^DKO^ mice. The major changes in BA were also reversed (Fig 3B). This is consistent with reduced hepatic expression in L^PPARαDKO^ mice of genes involved in cholesterol and BA synthesis (*Hmgcr*, *Baat*, *Cyp8b1 and Cyp27a*) and hepatocyte-mediated BA transport (*Abc11, Abc4, Abcg5, Abcg8)* (Fig 3C). Histological analyses indicated that PPARα deficiency increased liver steatosis, but reduced hallmarks of carcinogenesis (anisokaryosis, apoptosis, ductogenesis, dysplasia and mitosis) in L^PPARαDKO^ compared with L^DKO^ mice (Fig 3D). Moreover, CK19 and SOX9 staining were increased in L^DKO^ mice compared with L^PPARαDKO^ in agreement with cholangiocyte proliferation. Furthermore, although liver tumor lesions in L^DKO^ mice became prominently stained for glutamine synthetase (GS), reduced staining for glutamine synthetase was observed in the liver of L^PPARαDKO^ mice (Fig 3D). In addition, RT-qPCR analysis indicated that both inflammation and cholangiocarcinoma markers were reduced in L^PPARαDKO^ mice compared with L^DKO^ mice (Fig 3E, F). This evidence suggests that PPARα deficiency protected against the promotion of cholangiocyte proliferation in mice lacking hepatocyte JNK1/2. To evaluate whether PPARα deficiency and subsequent normalization of BA production blunted the FXR/FGF15/FGR4/ERK pathway, we evaluated FXR target gene expression. Hepatic RT-qPCR analysis indicated that *Fgf15*, *Shp* and *Fgr4* expression were reduced in L^PPARαDKO^ mice compared with L^DKO^ mice (Fig 3G). This is consistent with the observation of lower levels of ERK activation, detected by immunohistochemistry, in L^PPARαDKO^ cholangiocytes (Fig 3H).

**Figure 3.**
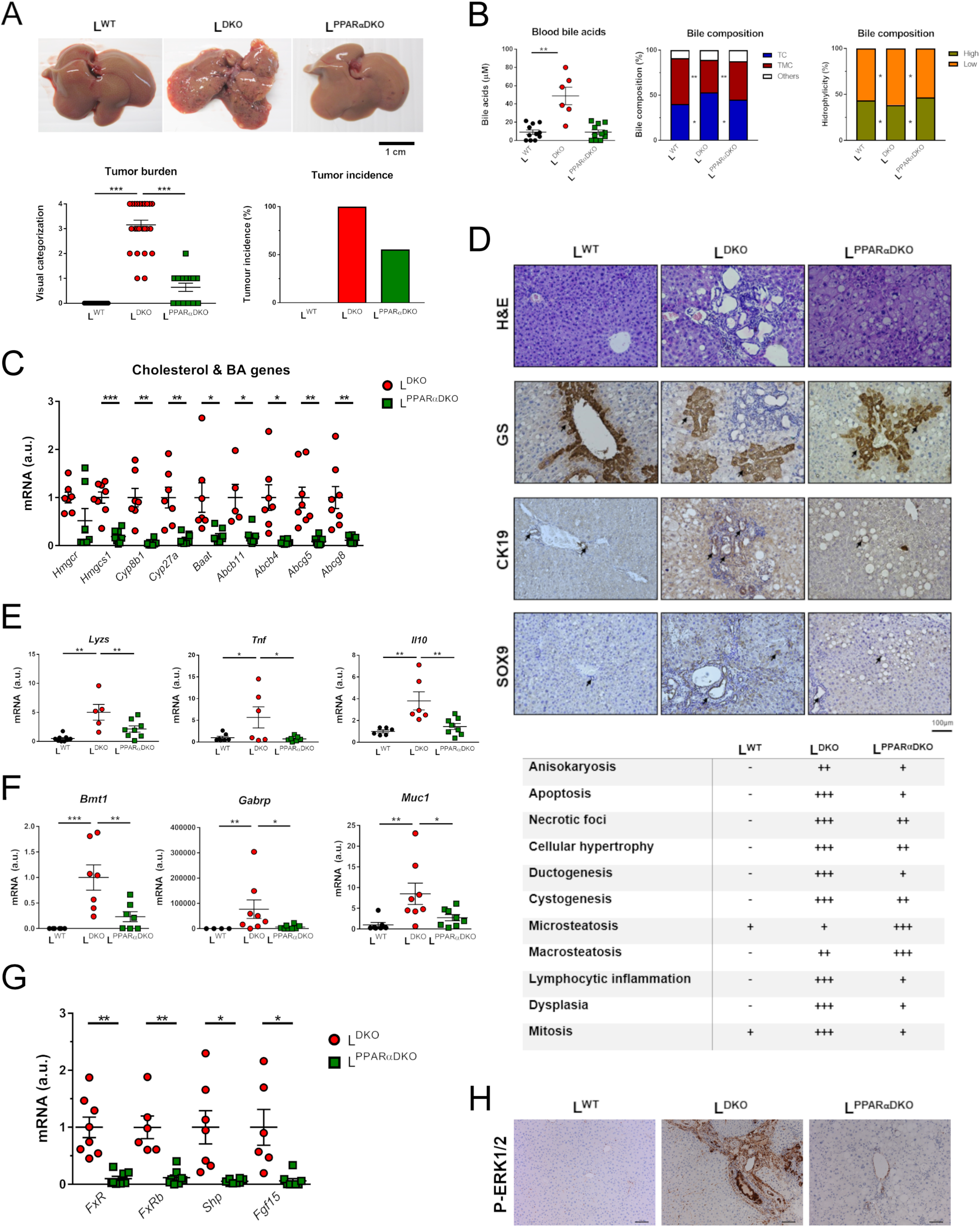
PPARα deficiency reduces liver cancer induced by hepatic JNK-deficiency. A Representative livers, tumor burden and incidence in 11 month old L^WT^, L^DKO^ and L^PPARαDKO^ mice (mean ± SEM; n = 14-25). B The amount of bile acid in the blood was measured (mean ± SE; n = 7-11). The composition of bile fluid collected from the gall bladder was examined. Statistically significant differences between L^PPARαDKO^, L^DKO^ and L^WT^ are indicated (*, P < 0.05; **, *P* < 0.01). C The expression of genes related to cholesterol synthesis (*Hmgcr* and *Hmgcs1*), bile synthesis and transporters (*Cyp27a1, Baat*, *Cyp27a*, *Cyp8b1, Abcb11, Abcb4, Abcg5, Abcg8*), were measured by quantitative RT-PCR (mean ± SEM; n = 5-8). The amount of mRNA was normalized to the amount of *Gapdh* mRNA in each sample. Statistically significant differences between LD^KO^ and L^PPARαDKO^ are indicated (*, P < 0.05; **, P < 0.01, ***, P < 0.001). D Representative liver sections of 10-month-old L^WT^, L^DKO^ and L^PPARαDKO^ mice stained with glutamine synthetase (GS), Cytokeratin 19 (CK19) and Sox9. Scale bar = 100 µm. E, F The expression of genes related to inflammation and cholangiocarcinoma was evaluated by quantitative RT-PCR. (mean ± SEM; n = 4-8). The expression was normalized to the amount of *Gapdh* mRNA in each sample. Statistically significant differences between LD^KO^ and L^PPARαDKO^ are indicated (*, P < 0.05; **, P < 0.01, ***, P < 0.001). G The expression of genes related to nuclear factor FXR pathway (*Fxr*, *Fxrb*, *Shp, Fgr4* and *Fgf15*) was evaluated in L^KO^ and L^PPARαDKO^ livers by quantitative RT-PCR. (mean ± SEM; n = 6-8). The amount of mRNA was normalized to the amount of *Actin* mRNA in each sample. Statistically significant differences between LD^KO^ and L^PPARαDKO^ are indicated (*, P < 0.05; **, P < 0.01). H Representative sections of the liver of 10-month-old L^DKO^, L^PPARαDKO^ and L^WT^ mice stained with Phospho-ERK. Scale bar = 100 µm.

### PPARα deficiency protects against DEN-induced HCC development in HFD-fed mice

Our analysis suggests that activation of PPARα may promote liver tumor development by altering BA physiology. Increased BA induces the synthesis and secretion of inflammatory cytokines in liver, which consequently results in liver injury (Miyake, Wang et al., 2000). Recently, it has been shown that increased hepatic BA controls HCC development in HFD-fed mice (Xie, Wang et al., 2016). To evaluate the role of PPARα in this context, we administered DEN to WT and PPARαKO mice on postnatal day 14, and 6 weeks later placed the animals on either a normal chow diet or a high-fat diet (HFD) in which 60% of calories are fat-derived (Park, Lee et al., 2010). Livers were examined for signs of HCC 8 months after DEN injection (Fig 4A). On normal chow diet, the two genotypes showed no significant differences in tumor number or size (Figu EV2). In contrast, on HFD, the mean number of tumors per animal was lower in PPARαKO mice compared with WT counterparts (Fig 4B). Moreover, tumors were smaller in PPARαKO mice than in WT mice (Fig 4B). This protection against HCC development in HFD-fed PPARαKO mice explained the better survival (Fig 4C). RT-qPCR analysis of tumor and non-tumor tissues from WT and PPARαKO mice revealed reduced expression of the cell-cycle genes *Cdk2*, *Ccna1*, *Foxm1*, and *Cdc25c* in non-tumor samples from PPARαKO mice and enhanced expression of the cell-cycle regulatory genes *p21*, *Trp53*, *p19* and *p57* in PPARαKO tumor tissue (Fig 4D). Collectively, these data indicate that the absence of PPARα protects against DEN-induced HCC in HFD-fed mice.

**Figure 4.**
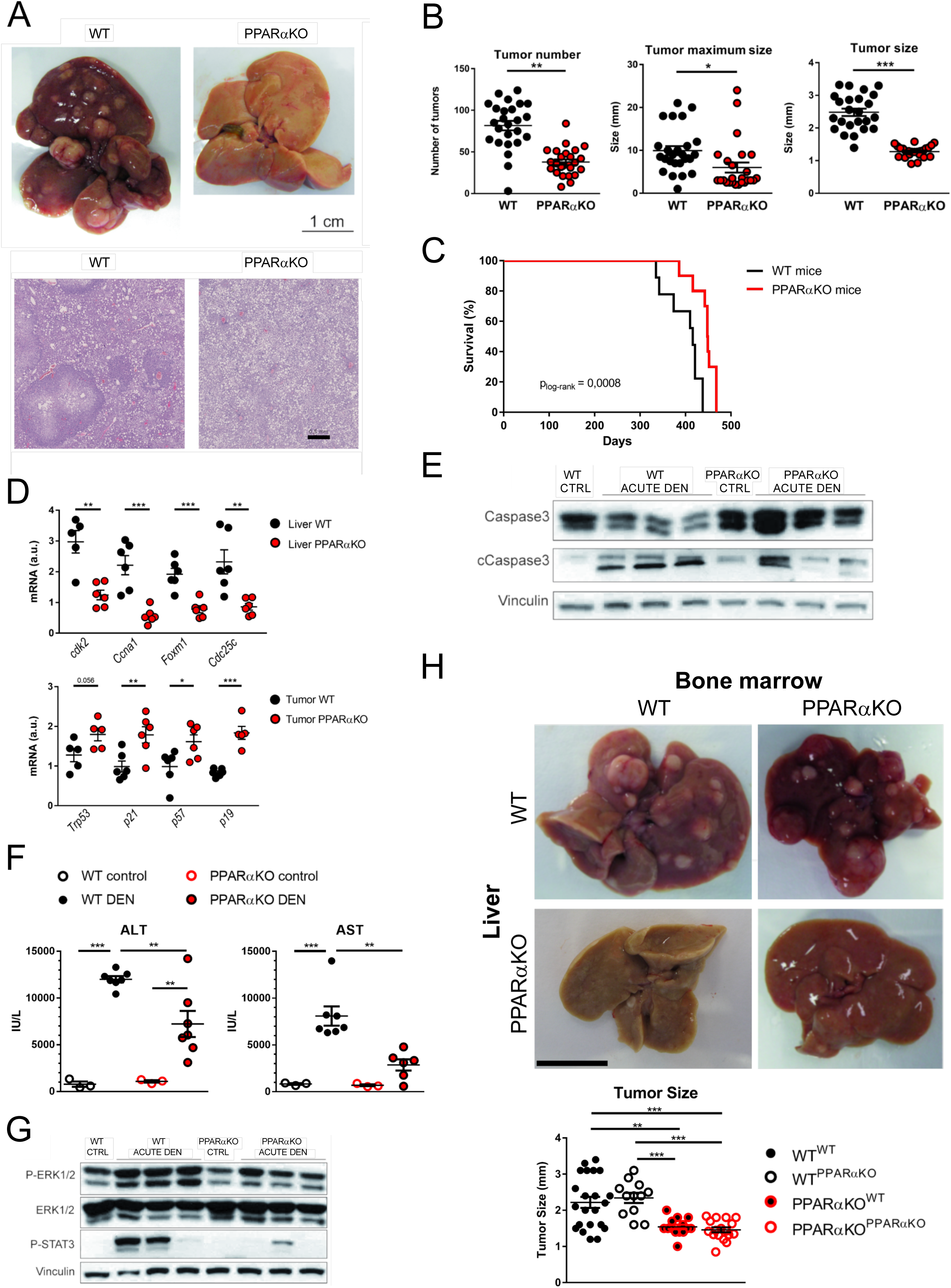
Effect of PPARα deficiency on HCC in HFD-fed animals. A Representative livers and H&E stained liver sections from diethylnitrosamine (DEN)-injected WT and PPARαKO mice fed a high fat diet (HFD) during 8 months. Scale bars = 1 cm / 0.5 mm. B Quantification of tumor number and size in HFD-fed DEN-injected WT and PPARαKO mice. The maximum diameter of individual tumor nodules (central panel) and the mean width of tumor nodules (right panel) are presented. (mean ± SEM; n = 25). Statistically significant differences between WT and L^PPARαKO^ are indicated (*, P < 0.05; **, P < 0.01, ***, P < 0.001). C Kaplan-Meier analysis of HFD-fed DEN-injected WT and PPARαKO mice (Mantel-Cox log-rank test; n = 9-10). D qRT-PCR analysis of cyclin and cell cycle regulator expression in liver samples from WT and PPARαKO mice. mRNA expression was normalized to *Gapdh* and WT liver expression (mean ± SEM, n = 5-6). Statistically significant differences between WT and PPARαKO mice are indicated (**, P < 0.01, ***, P < 0.001). E-G WT and PPARαKO mice were fed HFD from 6 weeks of age. At 19 weeks, mice were injected i.p. with DEN (100 mg/kg) and sacrificed 48 h later. E Immunoblot analysis of caspase3 and cleaved caspase3 in liver samples from untreated and acutely DEN-treated HFD-fed WT and PPARαKO mice. Vinculin protein expression was monitored as a loading control. F Liver damage was assessed from serum measurements of ALT and AST; (mean ± SEM; n = 3-7). Statistically significant differences between WT and PPARαKO mice are indicated (**, P < 0.01, ***, P < 0.001). G Immunoblot analysis of signaling pathways in liver samples from untreated and acutely DEN-treated WT and PPARαKO mice; blots were probed with antibodies to p-ERK, ERK, and p-STAT3. Vinculin expression was monitored as a loading control. H WT and PPARαKO mice were injected i.p. on postnatal day 14 with DEN (50 mg/kg). At 8 weeks of age mice were lethally irradiated and inoculated i.v. with bone marrow cells from WT or PPARαKO mice. After 2 weeks, mice were place on the HFD and then sacrificed at 8.5 months of age. DEN-induced liver cancers in WT and PPARαKO mice transplanted with WT or PPARαKO bone marrow (BM genotype is indicated as super index of recipient mice). The bar chart shows mean tumor size, and photographs show representative images of livers from each condition. (mean ± SEM; n = 12-22); Statistically significant differences between WT and PPARαKO mice are indicated (**, P < 0.01, ***, P < 0.001).

### PPARα deficiency protects from liver damage on HFD

DEN induces hepatocyte death associated with enhanced compensatory proliferation and augmented HCC development (Das, Garlick et al., 2011, Hui, Bakiri et al., 2007, Maeda, Kamata et al., 2005). DEN-induced apoptosis in liver tissue, measured by caspase 3 cleavage, was reduced in HFD-fed PPARαKO mice (Fig 4E). Moreover, analysis of ALT and AST levels induced by acute DEN treatment in HFD-fed mice revealed less liver damage in PPARαKO mice than in WT counterparts (Fig 4F). Additionally, ERK and STAT3, targets of FGF15/19 that modulate HCC development (Uriarte, Fernandez-Barrena et al., 2013, Uriarte, Latasa et al., 2015, Zhou, Luo et al., 2017), were less activated in livers of PPARαKO mice (Fig 4G).

However, there was no significant reduction in blood levels of the cytokines IL6, TNFα, IL1β, CCL2, and IFNγ in HFD-fed PPARαKO mice, suggesting that the lower acute DEN-induced liver damage was not associated with reduced inflammation (Fig EV3). After acute DEN injection, HFD-fed PPARαKO mice showed higher levels of the chemokines CXCL2 and CCL3 (Fig EV3), correlating with higher levels of markers of infiltration by immune cells (Fig EV4). These results suggest that PPARα in the liver, and not in hematopoietic cells, promotes HCC development in HFD-fed WT animals.

To confirm the role of PPARα in hepatocytes, we tested HCC development in chimeras created by transplanting WT or PPARαKO bone marrow (BM) into lethally irradiated WT or PPARαKO recipients. Chronic-DEN-induced HCC development was strongly suppressed in reconstituted PPARαKO mice compared with reconstituted WT mice, irrespective of donor BM genotype. On HFD, reconstituted PPARαKO mice also developed significantly smaller tumors than their WT counterparts, again irrespective of donor BM genotype (Fig 4H). Despite the established role of PPARα as an immune-modulator (Daynes & Jones, 2002), our bone marrow transplantation experiments show that loss of PPARα in immune cells does not contribute to the protection observed in PPARαKO mice. Our analysis indicates that that the protection against HCC in HFD-fed PPARαKO mice is not primarily mediated by bone-marrow-derived cells.

## DISCUSSION

The growing occurrence of liver cancer is due, in part, to an increasing prevalence of established risk factors, such as obesity and physical inactivity (Torre, Bray et al., 2015). HCC, in particular, is strongly associated with obesity and often appears after years of liver steatosis (Caldwell et al., 2004). In addition, long-term elevated BA levels are a risk factor for liver cancer development (Zhang, Zhou et al., 2015) and patients having elevated BA concentration and diabetes have a higher risk of developing HCC (Cui, Martin et al., 2018, Wu, Ge et al., 2010). Indeed, increased BA can lead to inflammation, apoptosis and necrosis of hepatocytes (Allen, Jaeschke et al., 2011, Jansen, Ghallab et al., 2017).

PPARα is an important modulator of liver metabolism controlling lipid and BA homeostasis, and its activation has been shown to decrease fatty liver disease (Abdelmegeed, Yoo et al., 2011, Li & Chiang, 2009, Yeon, Choi et al., 2004). We report that JNK mediated repression of PPARα causes changes in BA homeostasis which suppress cholangiocyte proliferation. Consequently, JNK-deficiency stimulates cholangiocyte proliferation and promotes the development of cholangiocarcinoma. This increased proliferation is mediated by the altered BA metabolism and the elevated hepatic expression of FXR/FGF15/FGR4 that triggers ERK activation in cholangiocytes. Our results have strong translational implications for obesity treatment. Activation of FXR by BA triggers the secretion of FGF15/FGF19 in humans (Inagaki, Choi et al., 2005), and the beneficial effects of FGF family on the obese metabolic profile has been well characterized (Nies, Sancar et al., 2015). However, their clinical use has been debated due to their implication in promoting liver cancer formation by stimulating proliferation (Cui et al., 2018, Zhou et al., 2017). In fact, it has been recently suggested that this deleterious effect of FGF15/FGF19 is more evident under HFD conditions (Cui et al., 2018). Our results support this observation as we describe a tumorigenic effect of PPARα under HFD condition and predict a deleterious effect of using FGF analogs for the treatment of obese patients.

Hepatic PPARα is an important mediator of this regulatory cascade. In fact, PPARα deficiency dramatically suppresses the phenotypes induced by JNK deficiency. The role of PPARα in liver cancer is still unclear. While some studies have demonstrated that PPARα activation might promote liver cancer (Hays, Rusyn et al., 2005, Nishimura, Dewa et al., 2007, Peters, Cattley et al., 1997), others indicate that PPARα activation may be neutral or suppress liver cancer development (Cheung, Akiyama et al., 2004, Heindryckx, Colle et al., 2009, Morimura, Cheung et al., 2006, Takashima, Ito et al., 2008). This could be due to different experimental conditions used in these studies. It is established that the consumption of a HFD causes PPARα activation (Soltis, Kennedy et al., 2017). The HFD also causes enhanced BA production and secretion which leads to cell damage and apoptosis, and the compensatory proliferation of surrounding cells (Sun, Beggs et al., 2016, Yoshitsugu, Kikuchi et al., 2019). Here we demonstrate that in WT mice PPARα promotes tumor development in a HFD DEN-induced model of liver cancer. These results contrast with the lack of effects of PPARα over tumor progression in WT mice fed with ND. The pro-tumorigenic effect of PPARα activation in WT mice is due, in part, to an alteration in BA metabolism that drives ERK activation. These conclusions are consistent with a recent report demonstrating that another nuclear receptor, PPARδ, is activated by HFD and promotes intestinal stem cells hyperproliferation driving to colon cancer (Beyaz, Mana et al., 2016). The data presented here identify obesity-induced PPARα activation as a critical factor in HCC development and progression. Moreover, our study provides evidence that in obesity the effect of PPARα is bone-marrow independent, and that a major inductor of liver cancer is the inhibition of the hepatic JNK.

Because of the role of JNK/PPARα/FGF signaling in lipid metabolism and carcinogenesis and their possible use for steatosis and obesity treatment, it is fundamental to understand the mechanism and conditions in which these signaling pathways might contribute to carcinogenic progression. JNK inhibition and PPARα activation are both potential therapeutic targets in obesity but our results urge to consider the risk of HCC development and other secondary effects during long-term treatments.

## MATERIALS & METHODS

### Animals

PPARα knock out mice (PPARαKO) (B6;129S4-Ppara^tm1Gonz^/J; RRID:IMSR_JAX:008154) and Albumin cre mice (B6.Cg-Speer6-ps1^Tg(Alb-cre)21Mng^/J; RRID:IMSR_JAX:003574) were purchased from the Jackson Laboratory and backcrossed for 10 generations to the C57BL/6J background (Jackson Laboratory; RRID:IMSR_JAX:000664). Mice with compound JNK1/2 deficiency in hepatocytes (L^DKO^) have been described (Das et al., 2011, Das, Sabio et al., 2009). Genotypes were identified by PCR analysis of genomic DNA isolated from mouse tails. All experiments were performed in male mice. For tumor studies, PPARαKO mice at postnatal day 14 received a single i.p. injection of 50 mg/kg body weight diethylnitrosamine (DEN, Sigma-Aldrich, N0258) dissolved in saline. Six weeks after DEN treatment, mice were put on a high-fat diet (HFD, Research Diet Inc.) or standard chow diet *ad libitum* until sacrifice 8 months after DEN injection. One group of HFD-fed mice was used for Kaplan-Meier analysis. For acute response studies, 6-week-old PPARαKO mice and WT mice were fed the HFD for 13 weeks, given a single 100 mg/kg body weight i.p. injection of DEN, and sacrificed after 48 hours. Radiation chimeras were generated by exposing 2-month-old DEN-injected recipient mice to 2 x 650 Gy ionizing radiation and reconstituting with 2×10^7^ cells from donor bone marrow by tail vein injection. Two weeks after bone marrow transplant, mice were fed the HFD *ad libitum* until sacrifice 8 months after DEN injection. Before sacrifice, blood samples were taken for analysis of ALT/AST and cytokines. In all cases, mice were euthanized after overnight starvation. Mice were housed in a pathogen-free animal facility and kept on a 12-hour light/dark cycle at constant temperature and humidity. All animal experiments conformed to EU Directive 2010/63EU and Recommendation 2007/526/EC, enforced in Spanish law under Real Decreto 53/2013 and the Institutional Animal Care and Use Committee (IACUC) of the University of Massachusetts Medical School.

### Serum analysis

Plasma activities of ALT and AST were assessed with the ALT and AST Reagent Kit (Biosystems Reagents) using a Benchmark Plus microplate spectrophotometer (Bio-Rad). Plasma concentration of non-sulfated bile acids were measured with the Bile Acid Assay Kit (Sigma-Aldrich) using a Fluoroskan Ascent fluorescence multiwell plate reader (Thermo Labsystems). Serum cytokine concentrations were measured by multiplexed ELISA (Millipore) with a Luminex 200 analyzer.

### Biochemical analysis

Total liver proteins were extracted in lysis buffer (50 mM Tris-HCl pH 7.5, 1 mM EGTA, 1 mM EDTA pH 8.0, 50 mM NaF, 1 mM sodium glycerophosphate, 5 mM pyrophosphate, 0.27 M sucrose, 1% Triton X-100, 0.1 mM PMSF, 0.1% 2-mercaptoethanol, 1 mM sodium ortovanadate, 1 µg/ml leupeptin, 1 µg/ml aprotinin). Extracts were separated by SDS– PAGE and transferred to 0.2 µm pore size nitrocellulose membranes (Bio-Rad). Blots were probed with primary antibodies to caspase-3 (#9662), cleaved caspase-3 (#9661), phospho ERK (#9101; RRID:AB_330744), ERK (#9102), phospho STAT3 (#9145; RRID:AB_2491009), and Vinculin (V4505, Sigma; RRID:AB_477617). All antibodies were used at 1:1000 dilution. After washes, membranes were incubated with an appropriate horseradish peroxidase-conjugated secondary antibody (GE Healthcare), and signal was detected using an enhanced chemiluminescent substrate for the detection of horseradish peroxidase (Clarity Western ECL substrate; Bio-Rad).

### Histochemistry

Histology was performed using tissue fixed in 10% formalin for 24h, dehydrated and embedded in paraffin. Sections (7 μm) were cut and stained using hematoxylin and eosin (American Master Tech Scientific). Sections were also incubated with Bouińs fluid overnight, counter-stain with hematoxylin (Sigma), and then stained with Masson-Trichrome stain (American Master Tech Scientific). Immunohistochemistry was performed by staining tissue sections with antibodies against PCNA (biotinylated from thermofisher MS-106-B; RRID:AB_64272), SOX9 (Abcam ab3697; RRID:AB_304012), glutamine synthetase (Abcam ab73593; RRID:AB_2247588), cytokeratin 19 (Abcam ab15463; RRID:AB_2281021) or phospho-p44/42 MAPK (Thr202/Tyr204) (Cell Signaling Technology #9101). Streptavidin-conjugated horseradish peroxidase (Biogenex), and the substrate 3,3’-diaminobenzidene (Vector Laboratories) were used followed by brief counter-staining with Mayer’s hematoxylin (Sigma).

### Analysis of biliary lipids

Bile was collected from the gall bladder following cholecystectomy. We determined cholesterol and phospholipids using an enzymatic assay (Wako). Total bile acids were measured using Hall’s Bile Stain Kit (American MasterTech). Bile acid species were examined by a modification of the method described by (Ye, Liu et al., 2007) using an HPLC-MS/MS (6410 Triple Quad LC/MS, Agilent Technologies). Chromatographic separation was achieved with gradient elution using a Zorbax Eclipse XDB-C18 column (150 mm x 4.6 mm, 5 µm) kept at 35°C and a flow rate of 500 µl/min. Initial mobile phase was 80:20 methanol/water, both containing 5 mM ammonium acetate and 0.01% formic acid, pH 4.6, and it was changed to 97:3 methanol/water over 9 min and then returned to 80:20 in 1 min. Electrospray ionization (ESI) in negative mode was used, with the following conditions: gas temperature 350°C, gas flow 8 l/min, nebulizer 10 psi, capillary voltage 2500 V. MS/MS acquisition was performed in multiple reaction monitoring (MRM) mode using the specific *m/z* transitions: [M-H]^-^ ion to 80,2 for taurine-conjugated bile acids and [M-H]^-^ ion to 74 for glycine-conjugated bile acids. Free bile acids did not generate characteristic ion fragments, as reported by others (Ye et al., 2007), and transition from un-fragmented precursor molecular ions 407.1 to 407.1, 391.3 to 391.3 and 375.3 to 375.3 were selected for trihydroxylated, dihydroxylated and monohydroxylated free bile acids, respectively.

### Real time q-PCR

Total RNA was isolated from liver and tumor tissue using the RNeasy Mini Kit (Qiagen) with on-column DNase I-digestion. Complementary DNA was synthesized with the High-Capacity Complementary DNA Reverse Transcription Kit (Applied Biosystems). Taqman^©^ assays were performed using the probes listed in Table 1 (Applied Biosystems). Sequences of primers used for quantitative real-time-polymerase chain reaction (qRT-PCR) are provided in Table 2. Expression levels were normalized to *Gapdh* and *Actb* mRNA. qRT-PCR was performed using the Fast SYBR Green system (Applied Biosystems) in a 7900HT Fast Real-time PCR thermal cycler (Applied Biosystems). A dissociation curve program was employed after each reaction to verify purity of the PCR products. The expression of mRNA was examined by quantitative PCR analysis using a 7500 Fast Real Time PCR machine.

**TABLE 1.**
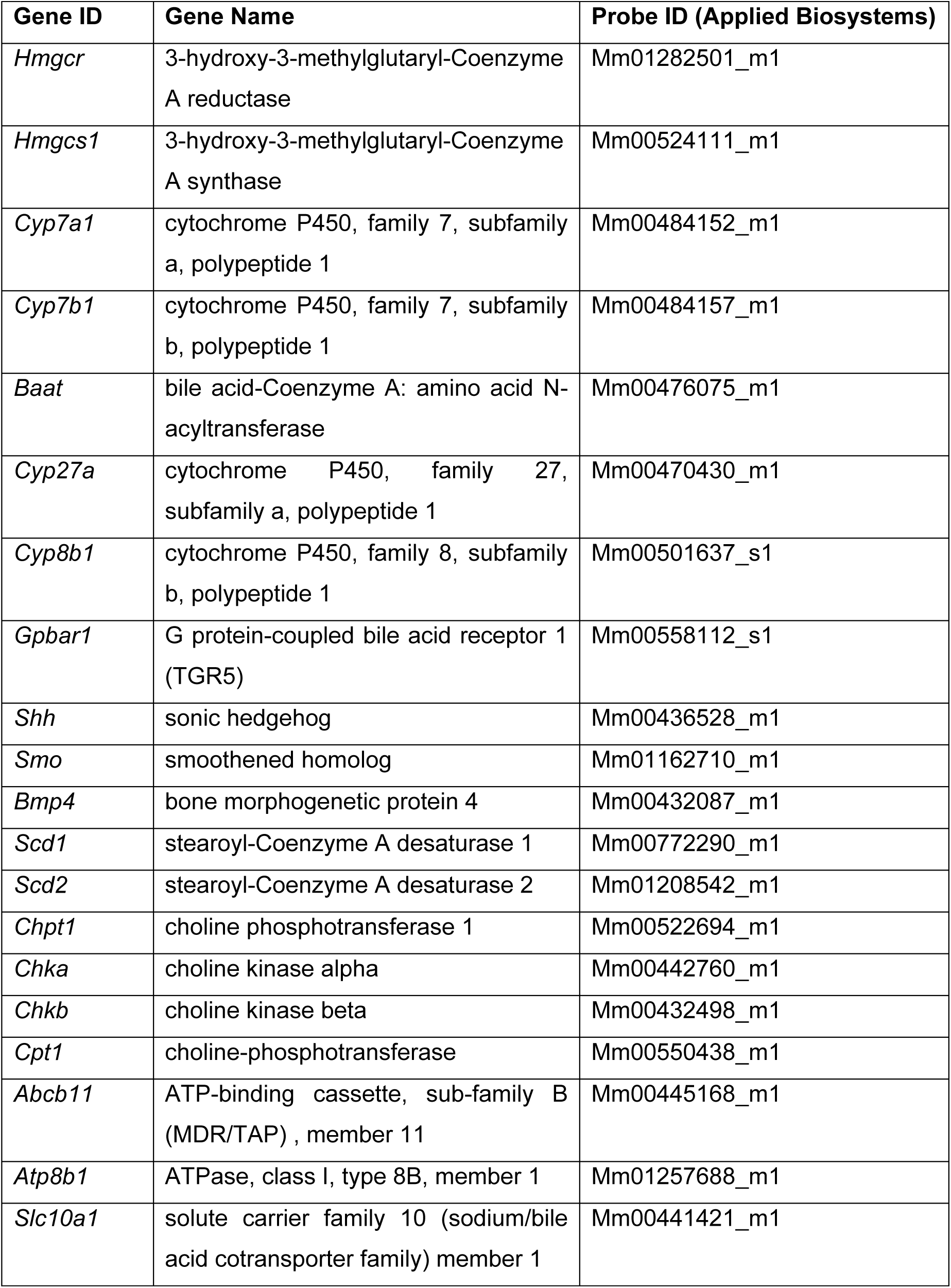
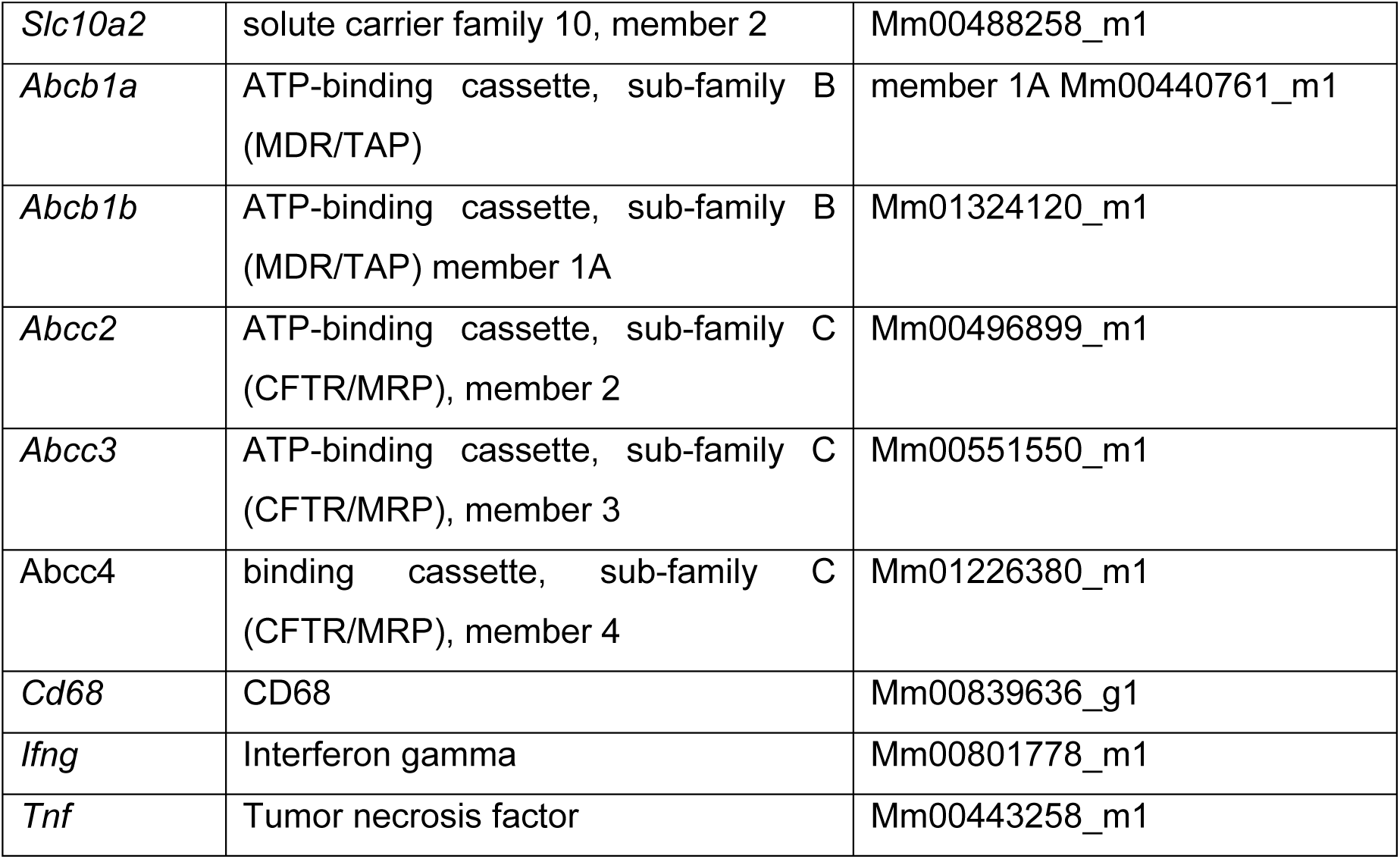
Taqman^©^ assays probes – Related to RT-qPCR

**TABLE 2.**
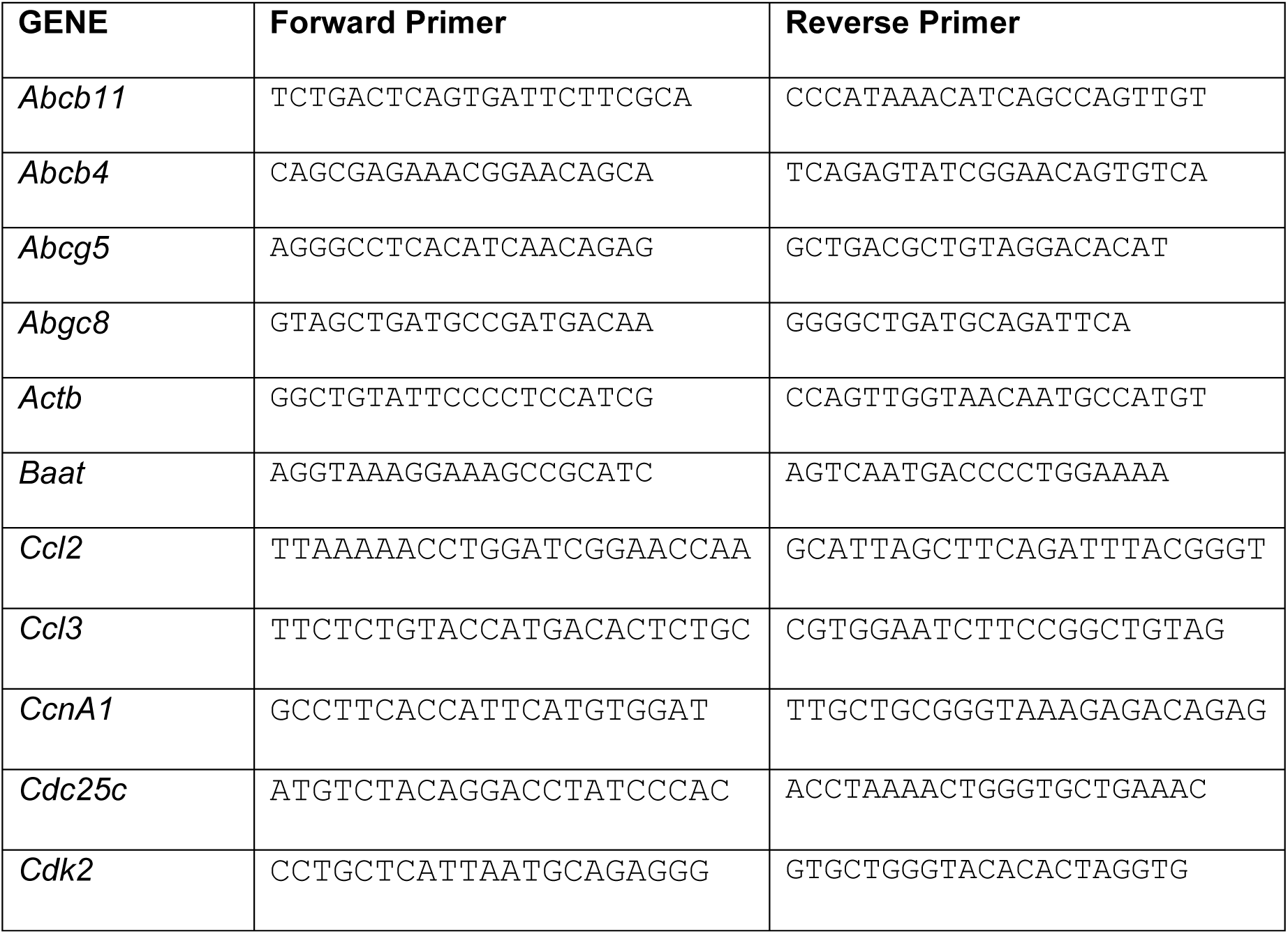
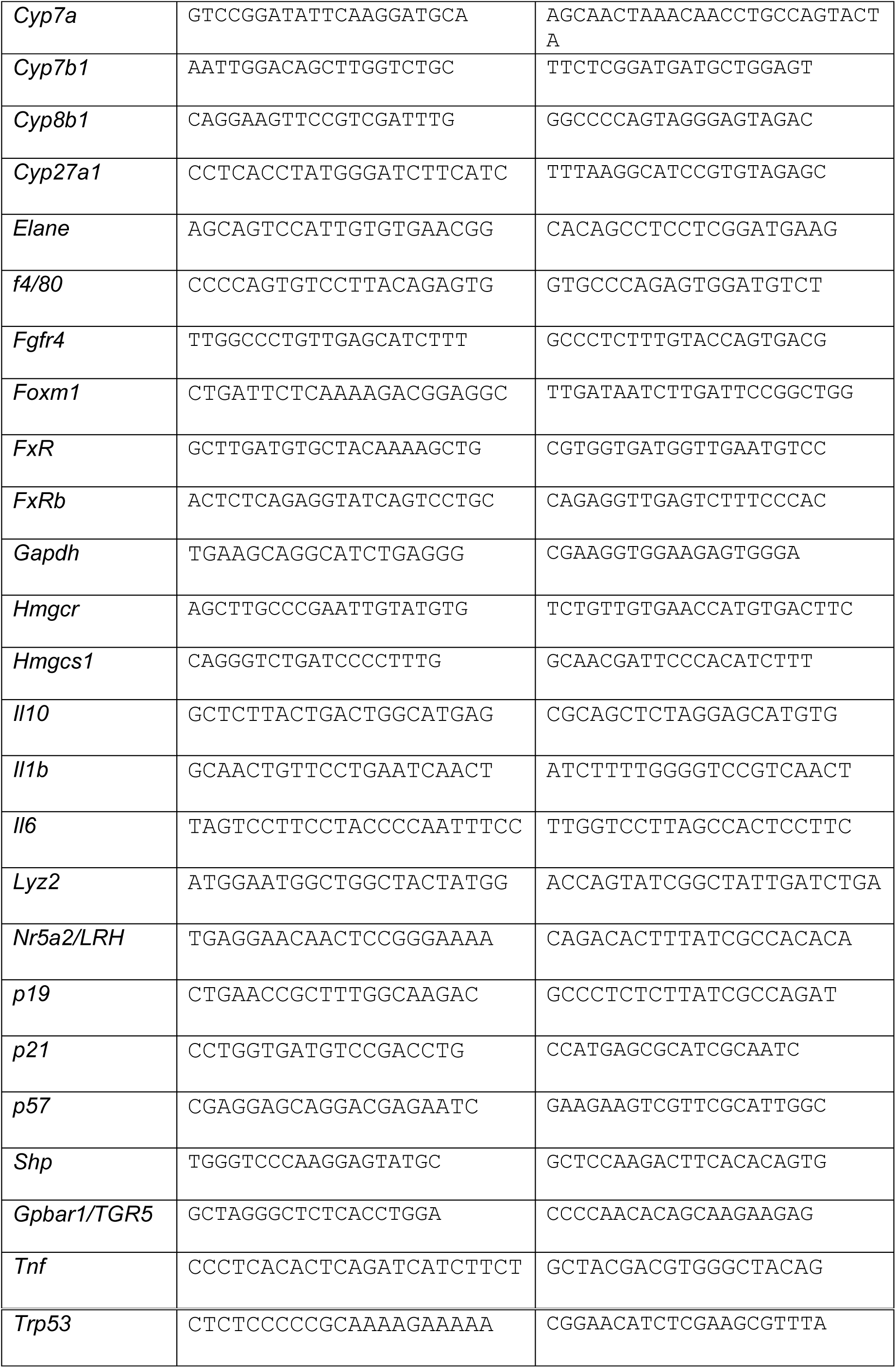
qPCR primers - Related to RT-qPCR

## QUANTIFICATION AND STATISTICAL ANALYSIS

### Statistical analysis

Differences between groups were examined for statistical significance using 2-tailed unpaired Student’s *t* test (with Welch’s correction when variances were different) or ANOVA coupled to Bonferroni’s post-test. Kaplan-Meier analysis was performed using the log-rank test. Statistical details and experimental n are specified in figure legends. Statistical analysis were performed with the GraphPad Prism 7 software (RRID:SCR_002798).

## ACKNOWLEDGMENTS

We thank S. Bartlett for English editing and David Garlick (University of Massachusetts Medical School) for pathological examination of tissue sections. CNIC Advanced Imaging and Animal facility for technical support.

G.S. (RYC-2009-04972), F.J.C (RYC-2014-15242) are investigators of the Ramón y Cajal Program. E.M was awarded La Caixa fellowship. This work was funded by grants supported in part by funds from European Regional Development Fund (ERDF): to G.S. European Union’s Seventh Framework Programme (FP7/2007-2013) ERC 260464, EFSD/Lilly European Diabetes Research Programme Dr Sabio, 2017 Leonardo Grant for Researchers and Cultural Creators, BBVA Foundation (Investigadores-BBVA-2017) IN[17]_BBM_BAS_0066, MINECO-FEDER SAF2016-79126-R, and Comunidad de Madrid IMMUNOTHERCAN-CM S2010/BMD-2326 and B2017/BMD-3733; F.J.C. EXOHEP-CM S2017/BMD-3727 and the COST Action CA17112.; F.J.C. MINECO Retos SAF2016-78711, the AMMF Cholangiocarcinoma Charity 2018/117, NanoLiver-CM Y2018/NMT-4949, UCM-25-2019, ERAB EA/18-14,. F.J.C. is a Gilead Liver Research Scholar. R.J.D: Grant DK R01 DK107220 from the National Institutes of Health; and to JJGM: PI16/00598 from Carlos III Institute of Health, Spain. The CNIC is supported by the Ministerio de Ciencia, Innovación y Universidades (MCNU) and the Pro CNIC Foundation, and is a Severo Ochoa Center of Excellence (SEV-2015-0505).

## Author Contributions

G.S. and A.M. conceived and supervised this project, E.M. and A.M. performed the experiments and prepared figures, G.S, A.M., R.J.D. and E.M. designed, developed the hypothesis, G.S, R.J. D, E.M, A.M., M J. M, J J.G. M analyzed and interpreted the data. L. E-L., T-L, M. E. R, D. S, C.C., F. J. C., L. L-V, T.B, J. C-K participated in the experiments. E.M., A.M and G.S. wrote the manuscript with input from all authors.

---

**Figure S1.**
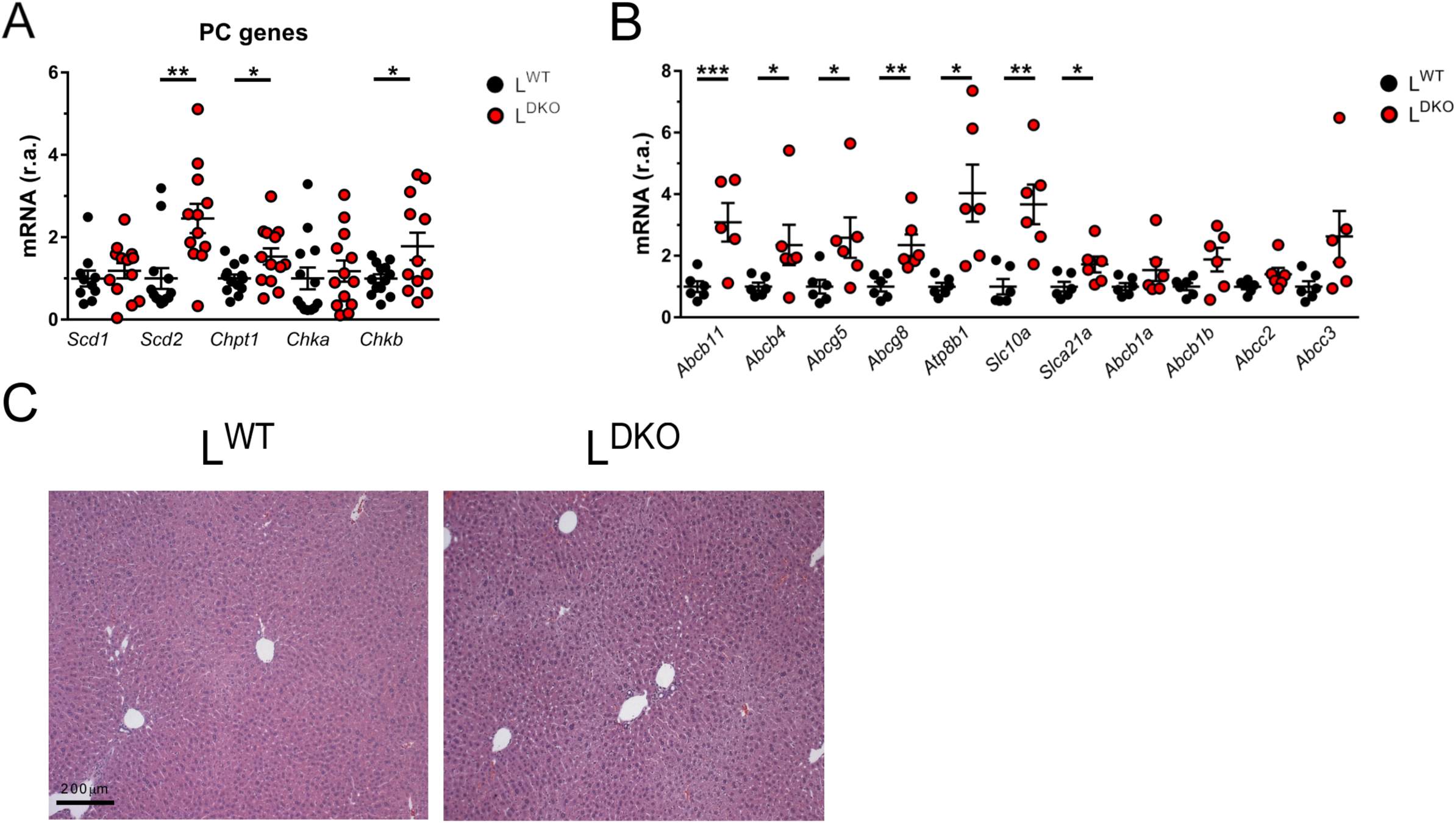
Effects of Hepatic JNK-deficiency liver. L^DKO^ and L^WT^ mice (age 4 months) were fasted overnight prior to removal of the liver. A, B The expression of genes related to phosphatidylcholine (PC) synthesis, hepatocyte-mediated transport of PC and BA was measured by quantitative RT-PCR. (mean ± SEM; n = 6-12). Gene expression was normalized to the amount of *18S* RNA in each sample. Statistically significant differences between L^DKO^ and L^WT^ are indicated (*, P < 0.05; **, P < 0.01). C Representative liver sections stained with hematoxylin and eosin (H&E). Scale bar = 200 µm.

**Figure EV2.**
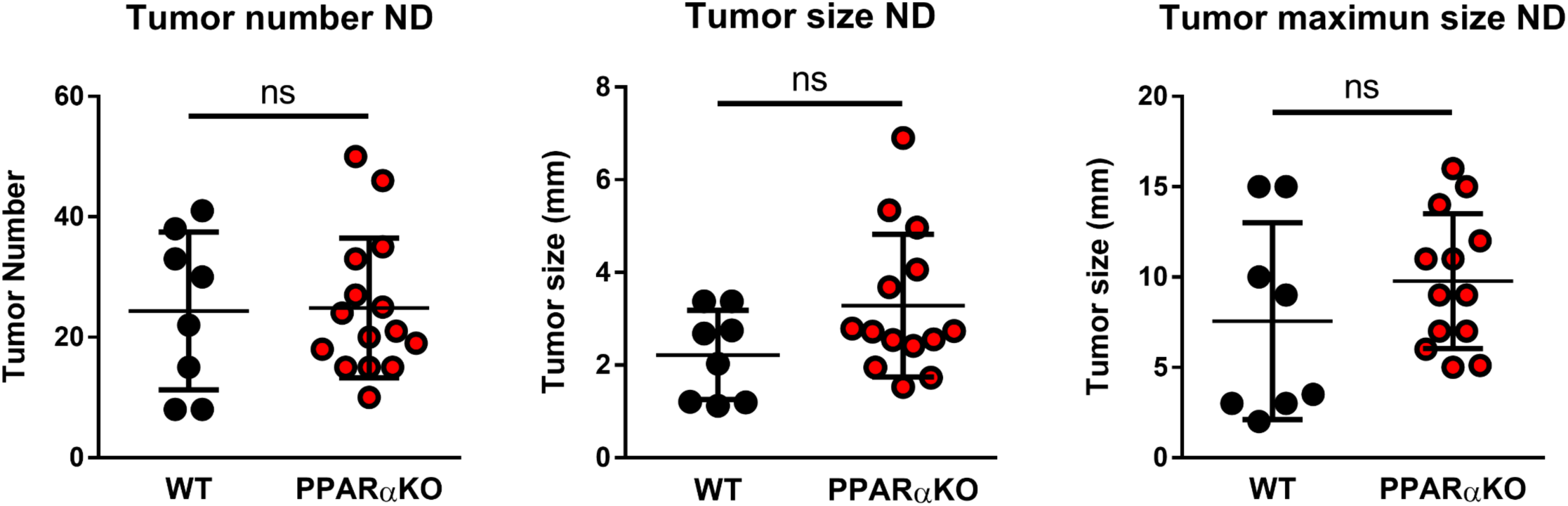
PPARα deficiency does not prevent HCC in mice fed a chow diet. WT and PPARαKO mice (14 day old) were injected with DEN and maintained on a standard chow diet (normal diet; ND). Tumor number and size were quantified after sacrifice 13 months post DEN injection. (mean ± SEM; n = 8-14). ns = No statistically significant difference.

**Figure EV3.**
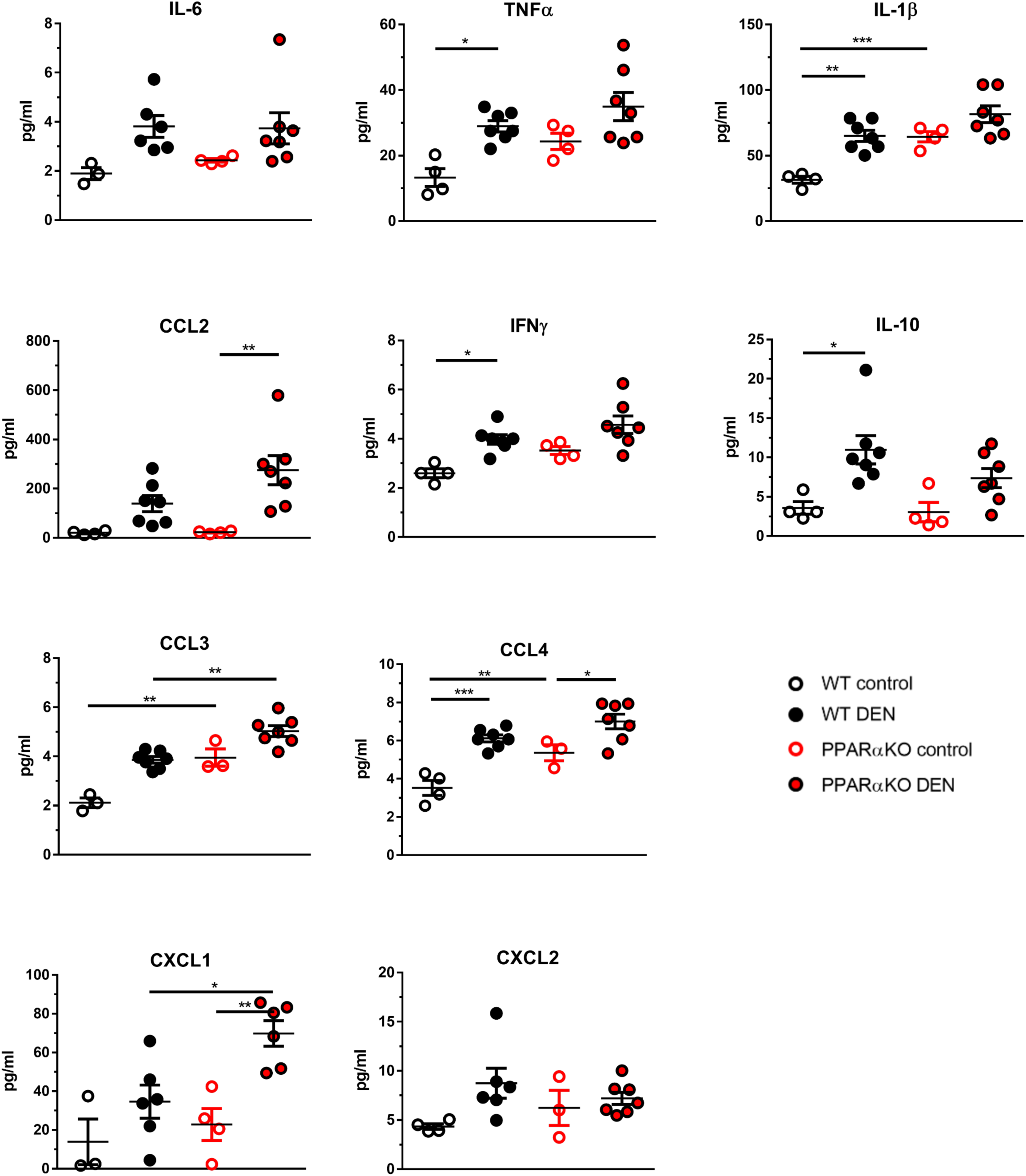
PPARα deficient mice on HFD exhibit defects in serum cytokines. Luminex analysis of cytokines and chemokines in blood from WT and PPARαKO mice. Mice were fed a HFD during 13 weeks and left untreated or acutely treated with DEN for 48 h (100 mg/kg). Circulating cytokines were measured. Data are shown as means ± SEM (n=4-7). Statistically significant differences between WT and PPARαKO mice are indicated (*, P < 0.05; **, P < 0.01, ***, P < 0.001).

**Figure EV4.**
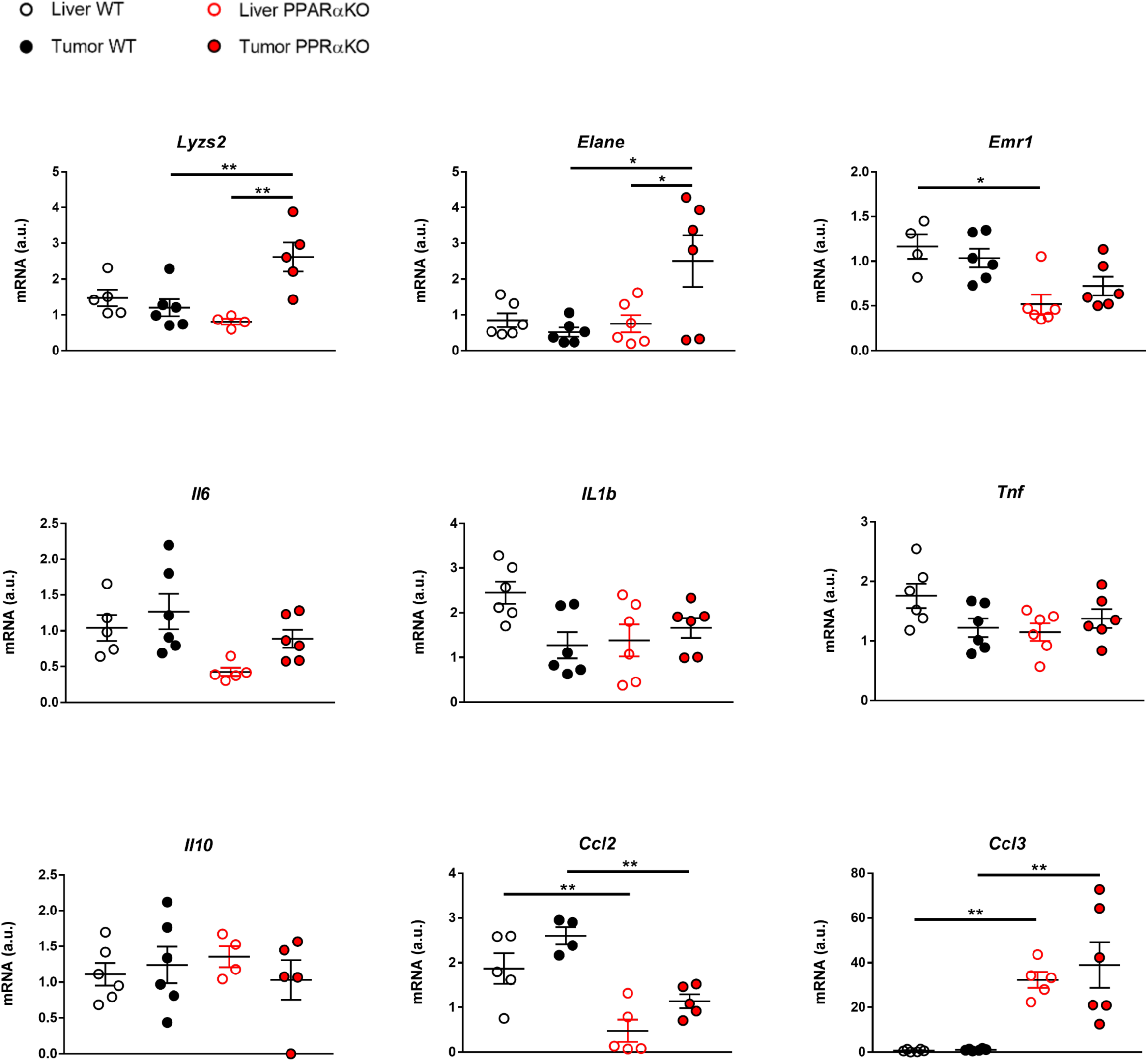
Effect of PPARα-deficiency on liver cytokine expression. RT-qPCR analysis of liver and hepatic tumor cytokine and chemokine expression in WT and PPARαKO mice. Mice were treated with DEN and maintained on the HFD for 8 months after DEN injection. The expression of genes coding for cytokines was measured by quantitative RT-PCR. mRNA expression was normalized to *Gapdh* and to WT liver. Data are shown as means ± SEM (n = 4-6). Statistically significant differences between WT and PPARαKO mice are indicated (*, P < 0.05; **, P < 0.01).

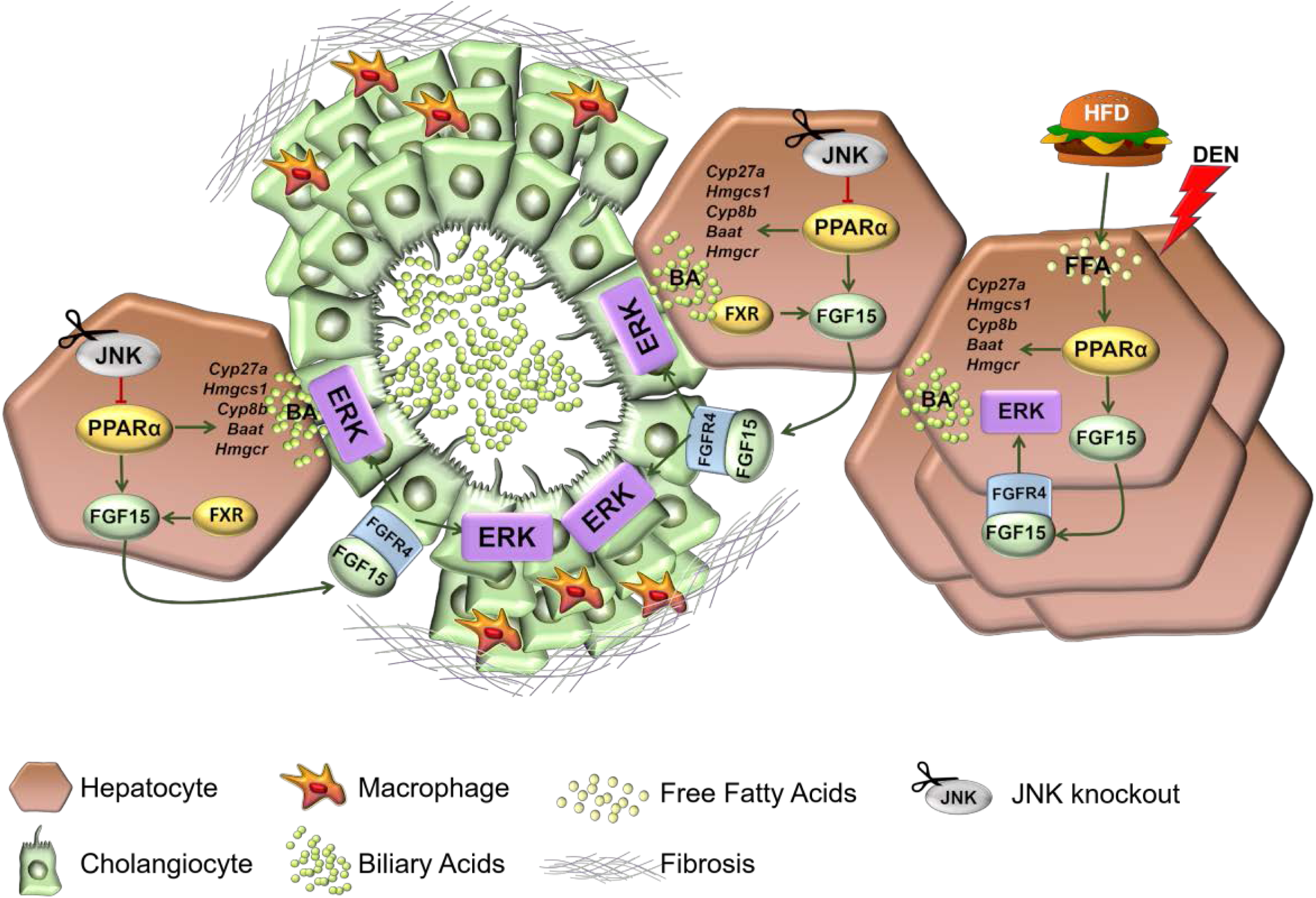

